# Spatio-temporal Characteristics of Noun and Verb Processing during Sentence Comprehension in the Brain

**DOI:** 10.1101/2020.06.22.163808

**Authors:** Sharmistha Jat, Erika J C Laing, Partha Talukdar, Tom Mitchell

**Affiliations:** Indian Institute of Science, Bangalore; School of Computer Science, Carnegie Mellon University

## Abstract

The human brain is very effective at integrating new words one by one into the composed representation of a sentence as it is read left-to-right. This raises the important question of what happens to the neural representations of words present earlier in the sentence? For example, do the strength of word representations encountered earlier on in the sentence remain constant or do they evolve as additional words are processed? Representation of words by neural activity in the brain has been the subject of several previous studies. We perform the experiment with a naturalistic task in which the subjects read simple active and passive sentences. Naturalistic studies have tended to explore words in isolation or in a very limited context (e.g., adjective-noun phrases). Representation of previously encountered words during incremental sentence reading, and how such representation evolve as more parts of a sentence are read, is a fundamental but unexplored problem – we take a first step in this direction. In particular, we examine the spatio-temporal characteristics of neural activity encoding nouns and verbs encountered in a sentence as it is read word-by-word. We use Magnetoencephalography (MEG) to passively observe neural activity, providing 1 ms temporal resolution.

Our experiments reveal that nouns and verbs read early in the sentence have a varying influence on neural activity while reading subsequent words, decreasing and increasing at particular word positions in active and passively voiced sentences, with particularly important contributions to activity in frontal and temporal cortical regions. We find the noun and verb information to be decodable from the neural activity for several seconds after sentence reading has completed. Our exploration is also the first to study the effect of question-answering task on the neural representation of the words post-sentence. We are releasing our 300 sentence MEG dataset to encourage further research in this important area.

## 1 Introduction

As we read a sentence word by word, our brains integrate the meaning of a newly encountered word to the composed representation of words read so far. This raises the question of what happens to the neural representation of words present earlier in the sentence. Are those words still actively retained in the composed representation? Or does the strength of their presence in the composed representation decrease over time? And what happens once the sentence has been read fully? These are fundamental questions whose answers are essential towards deepening our understanding of how sentences are processed in the brain. We initiate a study in this direction and provide empirical observations and insights for these questions. In particular, we use machine learning and naturalistic brain imaging experiments to study the spatio-temporal characteristics of noun and verb representations during sentence comprehension in the brain.

Semantic composition is the process of combining small units of linguistic information (words) to form meaning. Semantic composition has been the focus of several studies [1, 19, 20, 40]. A recent study [14] uses source reconstructed Magnetoencephalography (MEG) to investigate how individual words are interpreted in a sentence context by comparing the magnitude of brain activity between words in sentences and lists of words. Through their analysis, the study is able to detect the time at which the current word is combined into the current sentence context. However, some key questions like the composition of the current context are not answered. Machine Learning (ML) techniques can tell us more about the information content of a signal by testing the ability of that signal to distinguish between categories of interest. The application of ML techniques to naturalistic stimuli (as opposed to carefully chosen stimuli) enables us to study a wider range of scientific questions regarding the information content of the recorded brain activity. One naturalistic study successfully approached the question of semantic compositions in the restricted setting of adjective-noun phrase comprehension [7]. In this paper, we expand on that work to incorporate composed representations of words in a full sentence context with the aim of elucidating the contributions of the past and current words during the evolution of sentence-level meaning.

We also collect a new MEG dataset of 300 simple sentences in the active and passive voice. MEG [21] is especially ideal for understanding the fast dynamics of language processing due to its superior time resolution. Simple sentences allow us to focus on the brain processes involved in processing the words in a sentence, as opposed to the more complex syntactic elements in the sentence. We present these sentences word by word to the subject and to a BERT model [4]. Sentence representation by BERT model has been shown to be predictive of the brain activity data [16, 36]. We use the BERT representation vector as a state of the art representation of the composed meaning of the word sequence up to a given time point. A decoding framework is used to decompose the current context into constituent word representations at various time positions in the brain. The decoding accuracy is reported for the individual brain regions and the whole brain. In summary, we find the degree to which earlier words modulate the neural activity according to the BERT hypothesis about the composed meaning. Our experiments reveal important contributions from frontal and temporal regions of the brain in the mechanism of context composition during sentence comprehension. We also find the word representations to be decodable for a significant amount of time after the end of the sentence.

In summary, in this paper, we make the following contributions.

- We propose a new method to quantify the degree to which earlier words modulate the neural activity in the current context composition.
- Our results reveal important contributions from Frontal and Temporal regions of the brain in context composition. We also find the words to be decodable for a significant amount of time post-sentence.
- Our exploration is the first to study the effect of question-answering task on the neural representation of the words post-sentence.
- We also provide a new MEG dataset to study simple sentence understanding.

In the paper we outline the methods used to collect and process MEG data (Section 2), describe the framework to measure the predictive accuracy (Section 3), outline the setup and results of our experiments (Section 4), further discuss the related work (Section 5) and conclusions (Section 6).

## 2 SimplePassAct: A New Simple Sentence MEG Dataset

Simple sentences allow us to perform a focused study of word processing, as opposed to more complex syntactic elements in the sentence. Previously proposed MEG-based sentence datasets suffer from confounding issues, such as animacy and sentence length [26, 27]. To overcome these issues, we designed a new study with a larger 300 sentence dataset. We shall refer to the dataset as SimplePassAct dataset in the rest of the paper. Three participants (one female, two male) read 300 unique sentences, 150 active and 150 passive, comprised of six nouns (*singer, baker, customer, parent, artist, author*) and five verbs (*encouraged, attached, answered, challenged, followed*). To minimize uncontrolled semantic bias, the nouns were matched in gender neutrality [17] and both nouns and verbs were matched in frequency using the Corpus of Contemporary American English [3]. There were no repetitions of any sentence in this study, and the noun positions were completely counter balanced across the entire sentence set. Please find the stimulus sentences in the Appendix.

Neural activity was recorded using an Elekta Neuromag device (Elekta Oy). Sentences were presented across six blocks in randomized order unique to each participant. During each trial, each word of the sentence appeared on the display for 300ms followed by 200ms of blank screen, with 3.5-4s of rest before the next sentence began. To ensure participants engagement, 15% of the sentences were followed by a two-choice question regarding the agent or the patient of the preceding sentence and were given 2s to respond, followed by an additional 1.5s rest before the next sentence began. The two answer choices were presented. Left and right position of the correct answer was balanced across the experiment to eliminate perceptual and laterality bias. The MEG data was acquired adhering to the best practices of MEG data collection [10]. The collected data was spatially filtered using the temporal extension of SSS [35] which also realigned the head position to a default location. The Signal Space Projection (SSP) method [37] was used to remove artifacts captured by empty room data recorded on the day of each participant’s data acquisition, then the MEG signal was band-pass filtered from 1 to 150 Hz with notch filters applied at 60 and 120Hz to remove the contributions of line noise. The SSP method was again used to remove signal contamination by eye blinks or movements and cardiac signals.

The sentences are presented only once to our three subjects. This allowed us to present a higher number of sentences (300) in a given session. Though a group of three participants is on the smaller side for studies such as this, we note that the key features we describe in the results are apparent for most subjects and there is support for collecting more high-quality data from fewer subjects [18]. A study by Wehbe et al. [38] showed that the higher number of samples can be used by encoding models with good results. All three subjects consented to the study approved by the University of Pittsburgh and Carnegie Mellon Institutional Review Board.

## 3 Method

In this section, we present the methodological approach to explore semantic composition during simple sentence comprehension. We study words as they compose to form a context representation in the brain. The experiment is performed using naturalistic stimuli with simple sentences (Section 2). The sentence stimuli comprise of several similarly constructed sentences. This feature allows us to conduct experiments to compare representations among candidate sentences which differ by just one word. Some example sentences are, (a) Noun variation - “*the* ***artist*** *answered the parent*” v/s “*the* ***author*** *answered the parent*”, (b) Verb variation - “*the artist* ***answered*** *the parent*” v/s “*the artist* ***challenged*** *the parent*”. Figure 1 details the prediction framework. We predict brain activity using BERT representation (Section 3.1). The corresponding prediction rank accuracy calculation is described in Section 3.2. The candidate rank reference brain activity is computed as described in Section 3.3. Finally, the references are compared to the predicted brain activity to calculate a rank accuracy (Section 3.4).

**Figure 1:**
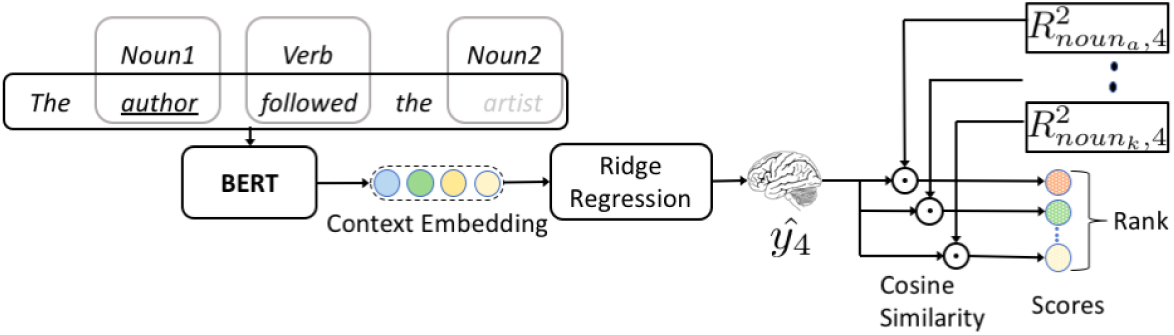
Decoding model for the MEG data. A string is processed by BERT model to obtain context embedding for the last token. For example, in the figure, the string “*the author followed the*”, is processed by BERT. The resulting context embedding is used to predict the corresponding brain activity using a learned Ridge Regression model (Refer Section 3.2 for more details). The candidate references are ordered based on the cosine similarity score with the predicted brain activity. The true candidate rank *r* is used to calculate rank accuracy. An above chance level (50%) accuracy suggests decodability of the stimulus. Please refer to Section 3 for more details.

### 3.1 BERT Representation

Bidirectional Encoder Representations from Transformers (BERT) is a state of the art Natural Language Understanding model [4]. The model is used to generate in-context embedding representations for text. The pre-trained model can be fine-tuned with additional layers to create models for a wide range of Natural Language Processing tasks. Unlike previous models, BERT can use both left and the right context to predict the randomly masked tokens. Jat et al. [16] show evidence of layer_18 representations to be the most predictive of brain activity. Similar to Jat et al. [16], we use layer_18 representations and process the sentences incrementally with BERT model to prevent information from future words from affecting the current representation, in line with how information is processed by the brain. For example, in the sentence *“the artist answered the parent”*, the representation of the word *“answered”* is calculated by processing sentence segment *“the artist answered”* and taking the last token representation from the layer 18, as the embedding representation for the word *“answered”*.

### 3.2 Decoding Model

We transform the neuroscience research question to a machine learning problem by learning a function 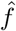 (Ridge Regression) to predict brain activity *y* from corresponding features of the stimulus *x* (substring representation). If the learned function 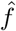 can accurately predict the brain activity data from the stimulus features for an unseen test example, then we conclude that the brain activity contains information about the stimulus. We measure the accuracy of the prediction using classification rank accuracy (Section 3.4). An above chance accuracy deems the stimulus decodable.

The brain activity data is preprocessed to improve the signal to noise ratio similar to Wehbe et al [39]. After the preprocessing, each trial *y* is represented using a (306, 500) sized matrix corresponding to the 306 sensors and 500ms time. The encoding model is learned to predict brain activity for 25 ms non-overlapping windows [23]. Each of the 25 ms brain activity is averaged over 5 ms non-overlapping windows to produce (306, 5) size brain activity *y*_*t*_ for the prediction task. We use Generalised cross-validation to train a ridge regression model [9] with regularisation parameter *λ ∈* [0.01, 0.1, 1, 10]. We compute the stimulus (substring) representation using BERT model Section 3.1. The predicted brain activity *ŷ*_*t*_ is evaluated based on a classification rank accuracy measure (Section 3.4). The prediction and classification experiment are run using cross-validation.

### 3.3 Candidate Reference Representation

Meaning of a word can become more specific as the sentence context unfolds, for example, the word “*artist*” can have different neural representation in the following two sentences “*the artist answered the parent*”, “*the artist challenged the parent*”. The brain as a perfect language machine updates its word representation in such a case. Therefore, it is appropriate to work with a candidate reference representation for neural representation of a word, which dynamically evolves with more sentence context. One simple way to compute this representation is by averaging all those sentences in which the word of interest appears in the same temporal position. But, for use in the rank accuracy computation, the candidates should be equally likely to occur in a test position. The following method computes these candidates with the desired properties.

We define the following notation to describe candidate reference representation. A single trial is represented by 500 ms of stimulus presentation time and is referenced as time position *i*. For example, in the sentence “*the artist answered the parent*”, the word “*artist*” is at time position *i* = 2 and “*parent*” at time position *i* = 5. The brain activity for a position *i* in sentence *s* is denoted by *y*_*s,i*_. A set of sentences where, word *w* is present at a fixed time position *α* in sentence *s*, is denoted by 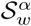. The word *w* can be represented with a candidate reference 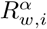 at any position *i ≥ α* in the sentence *s* as follows:

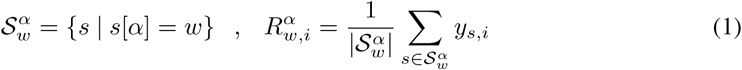

The rank reference 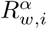 is the average brain activity of the word in a given context. When using these candidate reference representations in rank score computation, it’s important to make sure that each candidate for a given *α* position is equally likely. To make the candidates comparable, we add one more constraint on the set of sentences 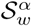. Any two pair of sentence sets *S* at given position *α*, should consist of same token-vocabulary (*V*_*S*_) except the candidate word at position *α* (Equation 2).

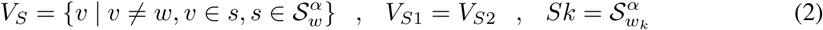

Please recall that our MEG dataset consists of sentences with 6 nouns, 5 verbs. These nouns and verbs are combined in all possible combinations to produce 300 active and passive sentences. Since a noun is not paired with itself in a sentence, the set of second nouns for any two given first nouns is distinct, thus violating the equal token-vocabulary condition (Equation 2). To satisfy the constraint, we divide the nouns *N* into two distinct sets as in Equation 3 to compute rank accuracy as follows:

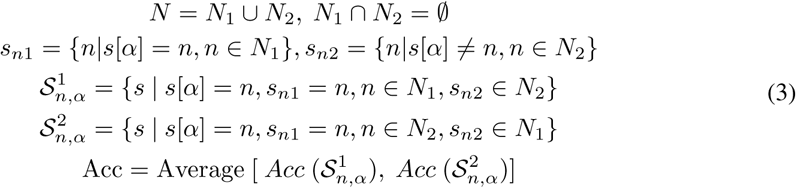

In short, the noun tokens are divided into two non-overlapping sets *N*_1_ and *N*_2_. The sentences are then grouped into a set 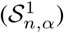, such that the noun1 is in *N*_1_ and noun2 is in *N*_2_ and vice versa for set 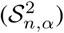. Rank accuracy is computed as the average decoding accuracy of the two sets 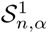 and 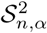.

### 3.4 Classification Rank Accuracy Measure

We use a classification rank accuracy measure to evaluate the presence of a noun/verb in the earlier sentence context. The rank accuracy measure [27] is more sensitive than the 0/1 measure commonly used in the literature [22]. This measure yields a confidence value (cosine similarity) over potential candidate references (Section 3.3). We order the references by the confidence values and assign a rank *r* for the correct candidate noun/verb from the set of candidates references *C*. The rank *r* is converted to an accuracy (Acc) using the Equation 4. The chance value for the rank accuracy measure is 0.5. Please refer to the Appendix for a proof.

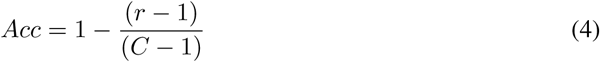

## Results

Experiments were performed with the SimplePassAct dataset (Section 2), over 25 ms non-overlapping time windows. The decoding accuracy is averaged over all the subjects. Due to the nature of our dataset, it is possible to encounter same text substring in both the train and test set, but with different brain activity data, for example “***the artist answered*** *the parent*” and “***the artist answered*** *the author*” have the same text substring till the third word. We note that this is not a strict leakage of information. However, to avoid any possible double-dipping into the test data, we performed careful cross-validation. For a noun condition, no two sentences with the same verb could separately occur in train and test split, resulting in a 5-fold (one fold for each verb) cross-validation. Similarly, for verb condition, a 6-fold cross-validation based on the first noun is performed. Please note that imposing the constraint in Equation 2 reduces the total number of sentences in the noun setting to 90 ([3 × 5 × 3] ×2). Both noun and verb are decoded from 0 ms to the end of the sentence and beyond. BERT representation of the last word in a sentence, is used to predict the brain activity for all end of sentence positions (eos1 – eos10 in Figure 3). While we do not expect the brain activity before stimulus onset to be able to decode the given stimuli, decoding accuracy at chance level works as a good sanity check for our results. The computations described in this paper were performed using a multiprocessor Intel Xeon E5 server with 377 GB RAM and 24 threads. The average run-time for computing a single experiment (for example, figure 2) is approximately 4 hours.

**Figure 2:**
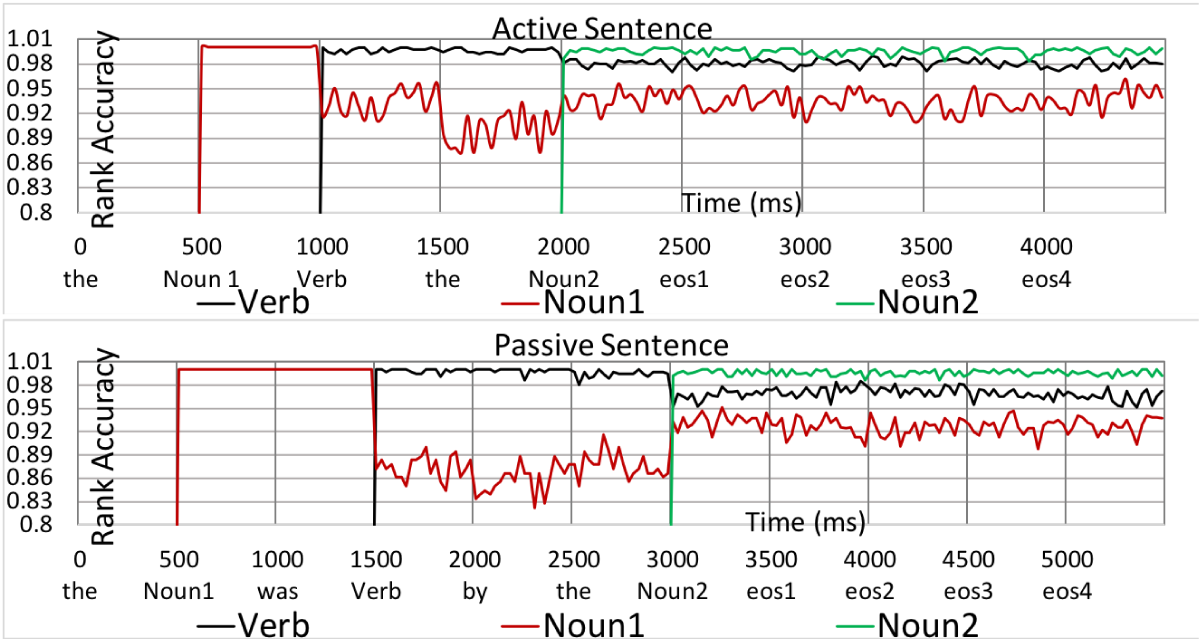
Classification rank accuracy of stimulus predicted from BERT layer-18 representations for noun1, verb and noun2 at various times during sentence comprehension. We continue testing post-sentence for an additional four-time positions (2000 ms). “eos” refers to “end of a sentence”. Our experiments reveal the resurgence of the first noun (noun1) representation when the second noun (noun2) is encountered in both active and passive sentences. And, that both noun and verb representations remain decodable after the sentence ends. Please refer to Section 4.1 for more details.

**Figure 3:**
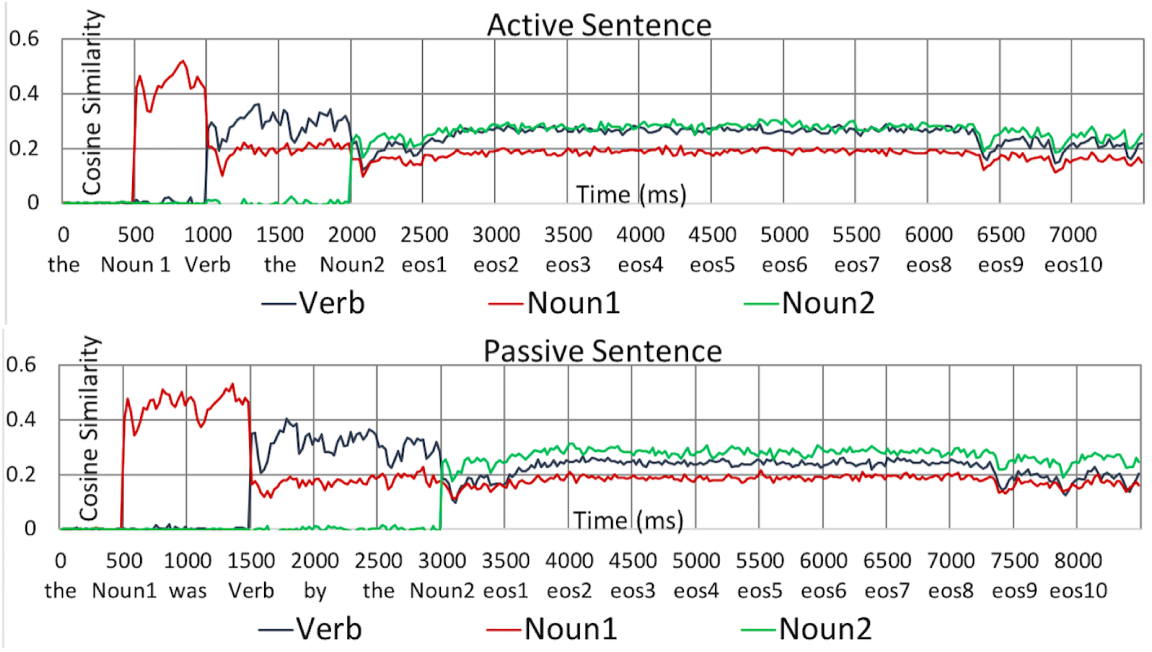
Cosine similarity of gold label data (irrespective of rank) predicted from BERT layer-18 representations for noun1, verb, noun2 at during sentence comprehension. “eos” refers to “end of a sentence”. We plot this information for an additional 5 seconds post-sentence. In the last 1 second (eos9, eos10) a new sentence is seen by the subject or a question (with 15% chance). We observe that the word representations are retained in the context until new information arrives when it slowly starts degrading. Please refer to Section 4.1 for more details.

### 4.1 Whole brain stimulus prediction accuracy

The rank accuracy of predicting noun1, verb and noun2 at various times during sentence comprehension is shown in Figure 2. We use the whole brain (306 sensors) data to predict the stimulus for this analysis. We find that the rank accuracy of all the words is high post-sentence. This makes sense, as the brain retains the representation of earlier sentence and therefore gets the “argmax(cosine similarity)” right in the rank accuracy measure. To test the degree of retention of this information, we also plot the cosine score for the gold-candidate in Figure 3. This plot displays accuracies till 5 seconds post-sentence, of which 4 seconds is a BLANK screen followed by 1s of next sentence stimuli or a question (with 15% chance). From the plot, we observe that the word representations are retained in the brain until a new piece of information arrives when it slowly starts degrading.

#### Noun Comprehension

We observe that the first noun (subject in active and object in passive) is decoded with significant accuracy during the word presentation and continues to be significantly decodable in the brain till the end of sentence activity. In addition, its decodability improves when a second noun is encountered and remains active even at 2s past the last word. Similar end of sentence decodability results was reported in Fyshe et al. [7]. The lack of predictive quality from noun1 to noun2 in the SimplePassAct dataset, might necessitate noun1 recollection for semantic composition when noun2 is shown.

#### Verb Comprehension

Similar to nouns, verbs are decodable with high accuracy during and after the presentation of the word. But, in contrast to nouns, the verb decodability remains heightened for the remainder of sentence reading. This result supports the growing body of literature for successful in-context verb decoding [16, 24].

#### Active v/s passive sentence comprehension

Both voices exhibit similar post-sentence neural representation. During the sentence reading, noun1 (object) in passive sentence displays different temporal decodability as compared to noun1 (subject) in active voice. We leave further exploration into this effect as future work.

### 4.2 When and where are the representations predicted in time?

The MEG helmet consists of 306 sensors, distributed over 102 locations. To understand the spatial distribution of word information in the brain, we predicted the noun1, verb, noun2 at different times with each sensor location (3 sensors) in the brain. The results from the analysis were then aggregated for larger brain regions (similar to Figure 1 in Hu et al. [13]). We examined the primary visual areas (left and right occipital lobe), speech and language processing areas (left temporal) and verbal memory (right temporal), sensory perception (left parietal) and integration (right parietal), language related movements (left frontal) and non-verbal functioning (right frontal). The aggregated brain region accuracy is shown in Figure 6 for verb stimuli. The left, right frontal and right temporal brain regions are the most accurate at predicting word representation. These brain regions have been implicated in memory and executive function in language networks [29, 5]. We find similar trends for nouns, please find more information in the Appendix section.

**Figure 4:**
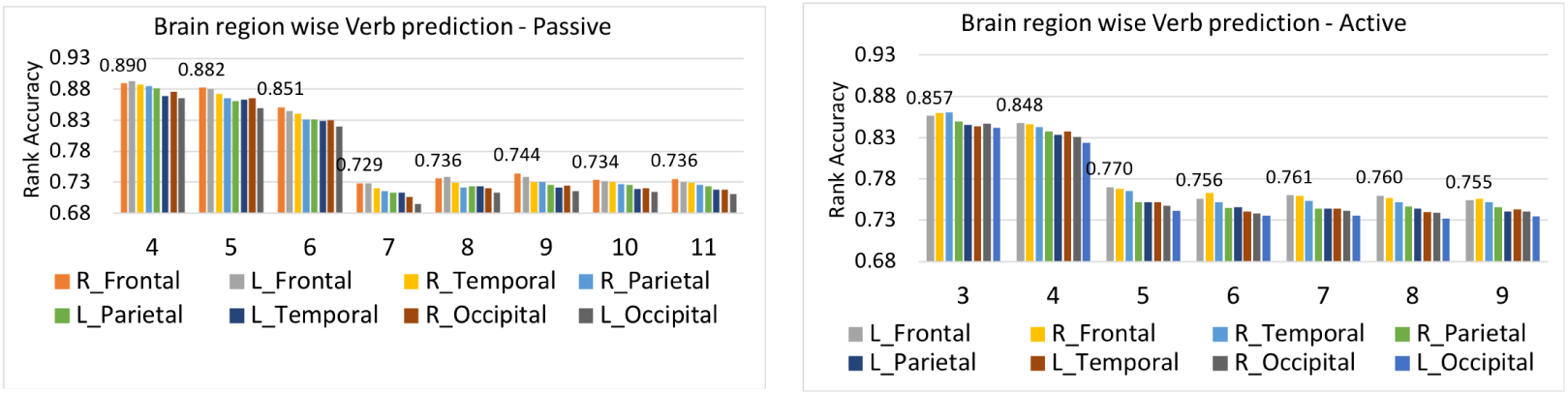
Rank Accuracy of predicting stimulus from each sensor location (102 locations each with 3 sensors). The resulting rank accuracy was aggregated in signal space over brain regions delineated by the position of each sensor in the MEG helmet. The x-axis represents the word position in the sentence. Each word position is 500 ms of brain activity data. The figure shows verb decoding accuracy from the position of occurrence to the end-of-sentence activity. We find that left, right frontal and right temporal regions are the most accurate at predicting word representation during sentence comprehension. These Brain areas underlying our outlined sensor regions have been implicated in memory and executive function by previous neuroscience research. Please refer to Section 4.2 for more details.

**Figure 5:**
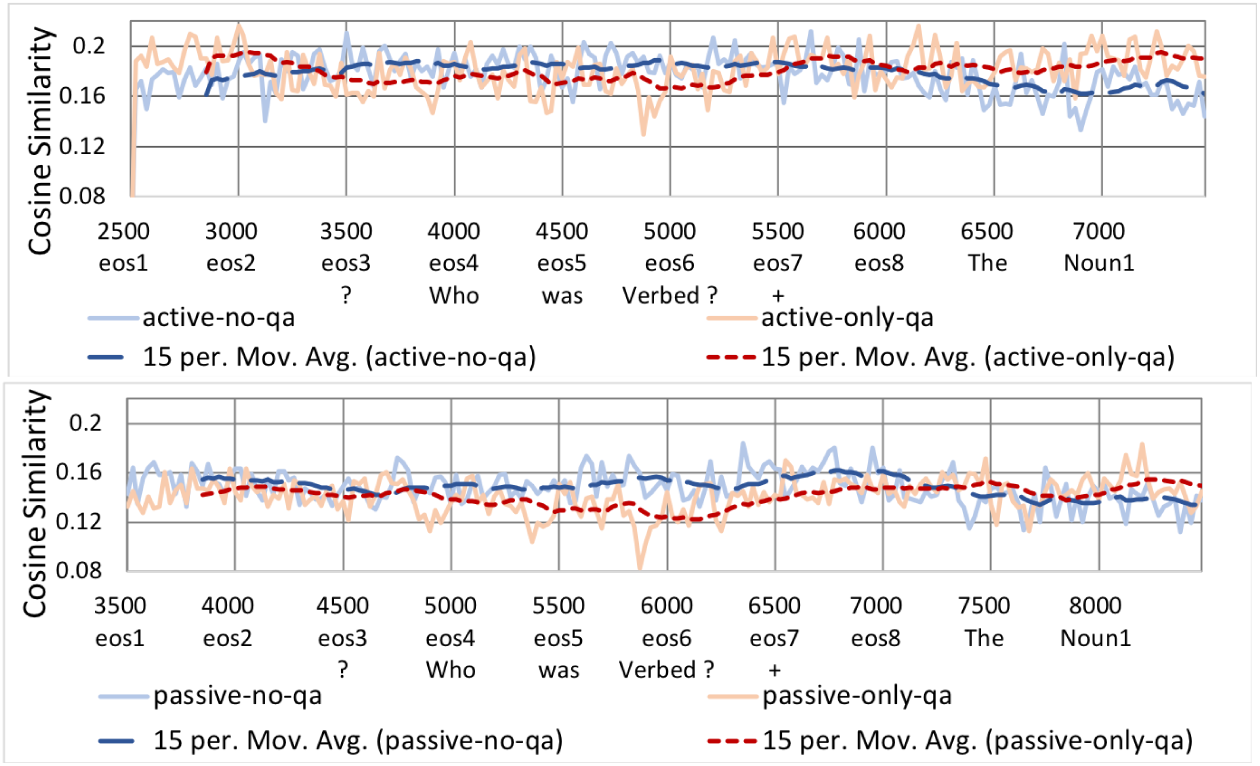
Comparison between post-sentence representations in “Only-QA” versus “No-QA” set, and the 15 point moving average of the corresponding series. We observe that, in the case of a question, the subject retains the sentence information for a longer period of time as compared to the no question set. Please refer to Section 4.3 for more details.

**Figure 6:**
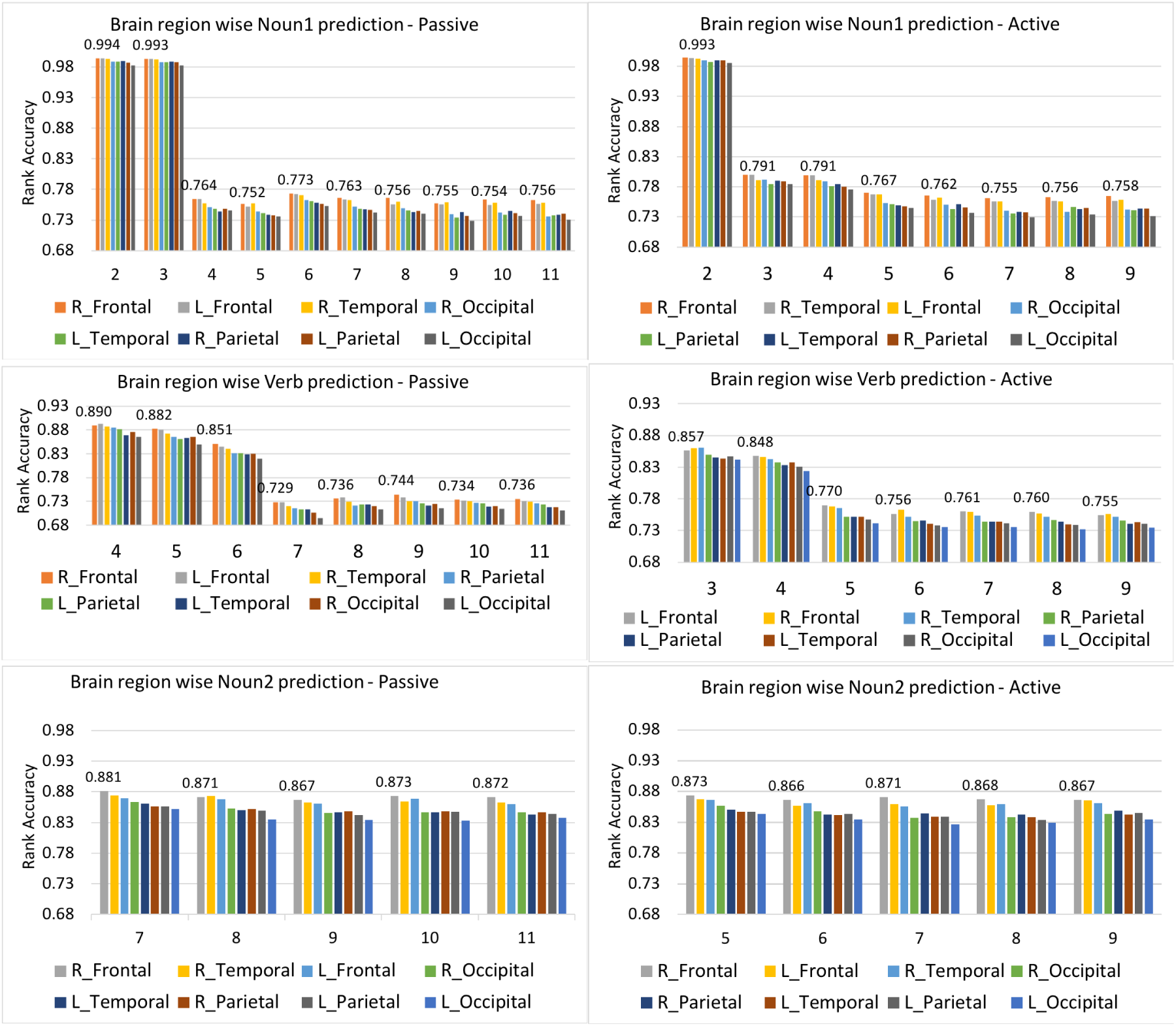
Rank Accuracy of predicting stimulus (noun1, verb, noun2) from each sensor location (102 locations each with 3 sensors). The resulting rank accuracy was aggregated in signal space over brain regions delineated by the position of each sensor in the MEG helmet. The x-axis represents the word position in the sentence. Each word position is 500 ms of brain activity data. For example noun1 appears at position 2 in the active sentence, the figure shows noun1 decoding accuracy from position 2 to end-of-sentence activity. We find that left, right frontal and right temporal regions are the most accurate at predicting word representation during sentence comprehension. Please refer to Section 4.2 for more details.

**Figure 7:**
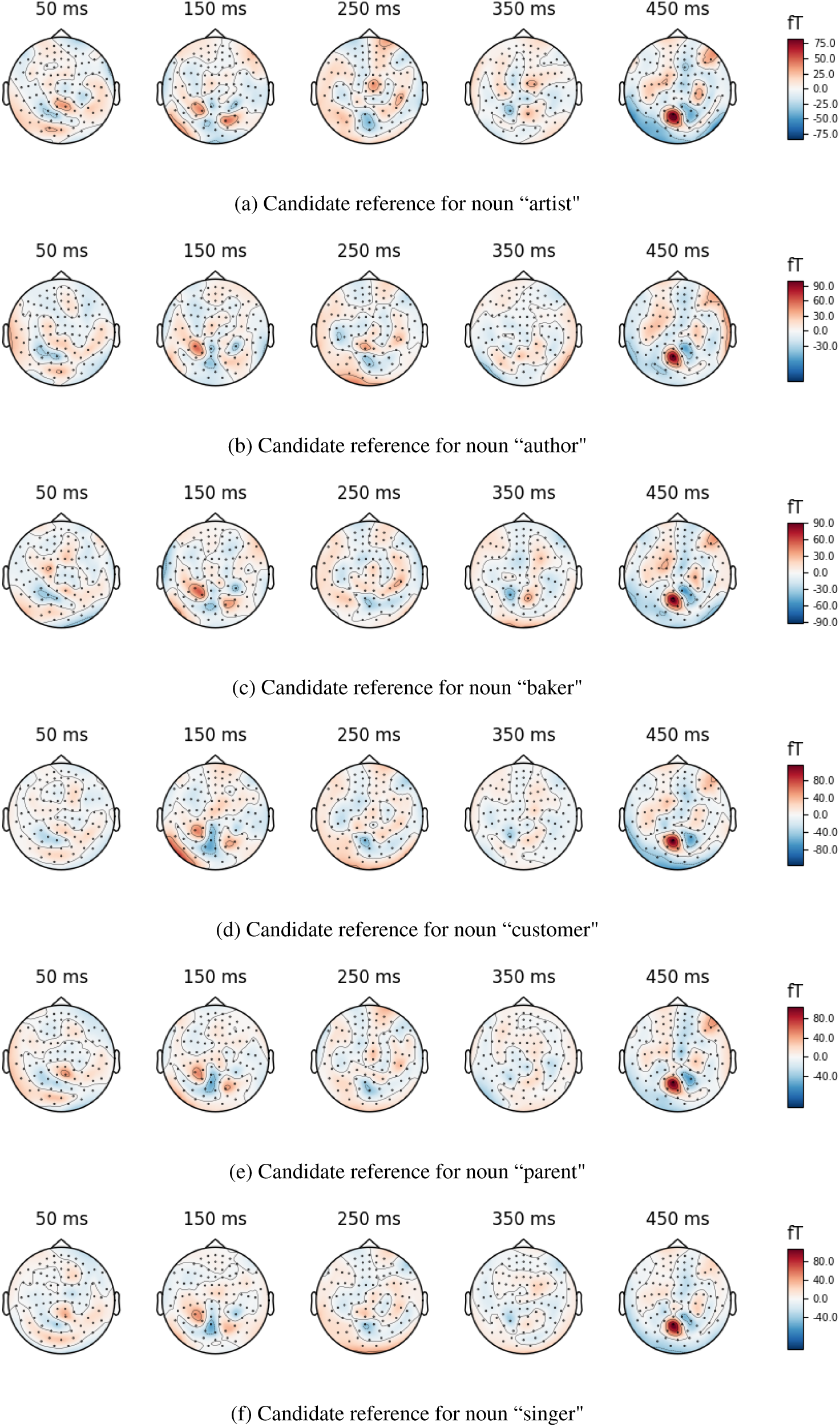
Example topomap visualisation of the candidate references for noun1 at word position (*α*) 4 in an active sentence (eg. “the artist answered the”). The brain activity data is shown for 500 ms with 100 ms non-overlapping windows. We observe distinct activity patterns in the candidate references from the visualisation. These differences are captured by the ML algorithms to help predict the correct stimulus. Please refer to Section 3.3 for more details.

**Figure 8:**
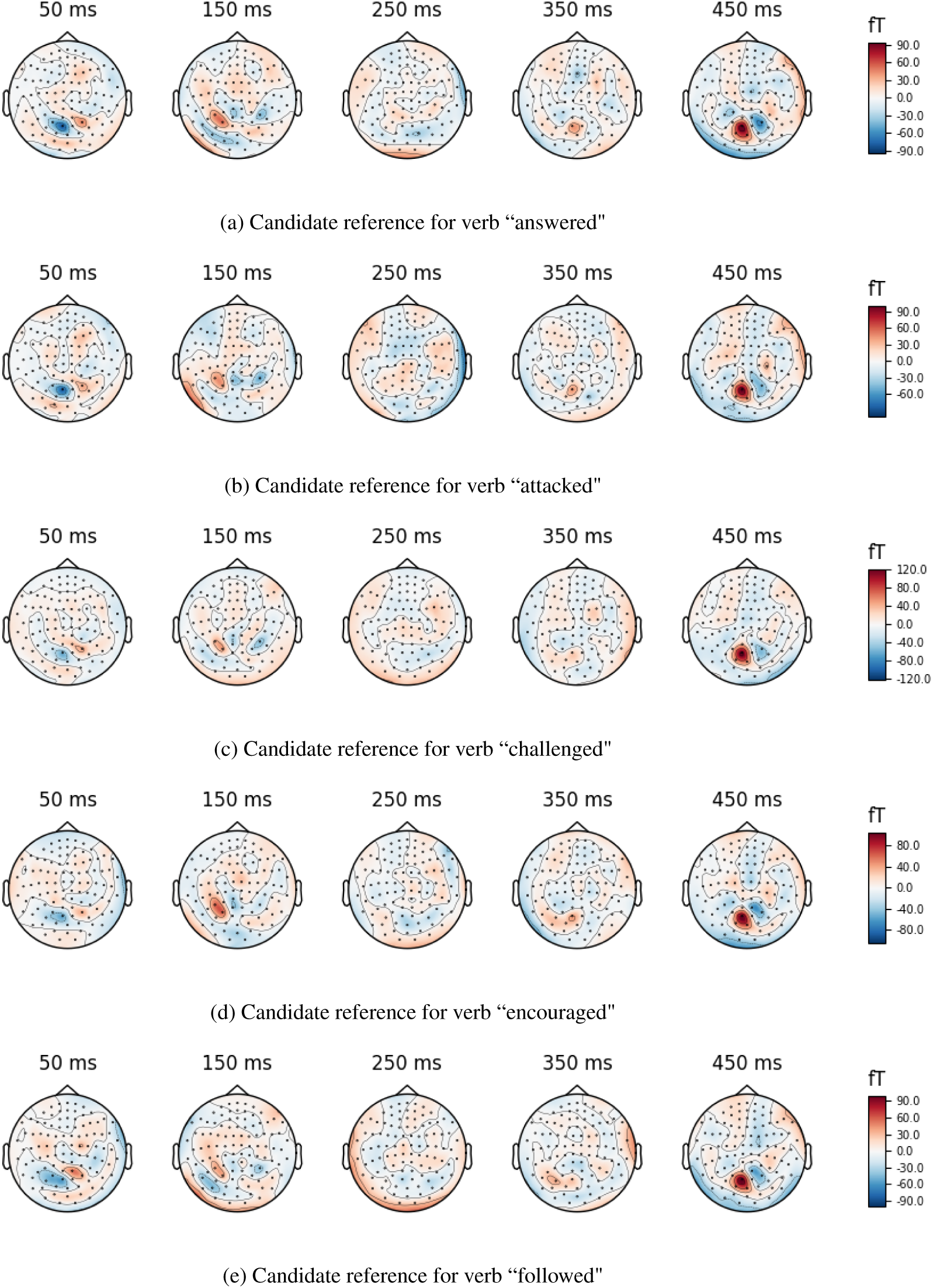
Topomap visualisation of the candidate reference for verbs at word position (*α*) 6 in a passive sentence (eg. “the artist was answered by the”). The brain activity data is shown for 500 ms with 100 ms non-overlapping windows. We observe distinct activity patterns in the candidate references from the visualisation. Please refer to Section 3.3 for more details.

### 4.3 What happens when a question is encountered?

In SimplePassAct dataset, 15% (45) of the sentences are followed by a question and then a short pause to allow the user to answer that question. How do the neural representations of the words presented earlier in the sentence evolve when a question is encountered? In Figure 5, we show the comparison of cosine similarity scores of earlier Noun1, Verb and Noun2 between conditions “during a question presentation” (Only-QA), vs “during blank screen before another sentence” (No-QA). Please note that the subjects do not know which sentences are followed by a question. An upcoming question is indicated by “?” sign, which appears on the screen between 800-1000ms after end of a sentence. To make the “Only-QA” and “No-QA” setting comparable in terms of the regression model, we sub-sampled the sentences in the “No-QA” setting to match that of “Only-QA”, approximately 23 sentences in each sentence voice. We observe, in the case of a question, the subject retains the sentence information for a longer period of time as compared to the no question setting, perhaps to answer the question. All three subjects had close to 100% accuracy in correctly answering these engagement questions.

## 5 Related Work

To study human language comprehension, researchers have used neuroimaging methods such as Functional Magnetic Resonance Imaging (FMRI) [8], electroencephelography (EEG) [2] and Magnetoencephalography (MEG) [21] etc. Evidence from deficit studies, FMRI, MEG, Event related potential (ERP) imaging studies have been combined to formulate neural models of language comprehension [6, 11, 12, 28]. Recently, Machine learning (ML) models are becoming popular in understanding the brain imaging data in the field of neuroscience. These models can be very sensitive in determining what stimulus information is present in the brain activity data. In addition, Deep Learning models trained on a large corpus of data are also used to provide rich stimulus features for the ML models [15, 39, 30]. Prior research by multiple papers has established a general correspondence between a computational model and the brain’s response to naturalistic language [22, 33, 25, 34, 32, 31]. Some studies have successfully used the BERT model to predict the brain activity data [16, 36]. For sentence comprehension, event related potential (ERP) studies present anomalous sentences to the subjects to study the timing (P600, N400) of various composition effects. In contrast, few studies understand the semantic composition using naturalistic methods, specifically methods that include stimuli with no surprisal effect or semantically predictable quality [7]. We add to these previous research works to further our understanding of sentence comprehension.

## 6 Conclusion

In this paper, we present a framework to study how neural representations of words present earlier in a sentence evolve as subjects read a sentence from left-to-right. More precisely, we focus on the representations of noun and verb in the neural encoding of a sentence. Our experiments reveal the resurgence of the first noun (noun1) representation when the second noun (noun2) is encountered in both active and passive sentences. The verb representation remains important from the onset of its presentation through the remainder of the sentence. Both noun and verb representations remain decodable after the sentence ends. We also reveal important contributions from Frontal and Temporal regions of the brain in context composition. Our exploration is the first to study the effect of question-answering task on the neural representation of the words post-sentence. We find that post-sentence question-answering helps in maintaining sentence representation in the brain. Finally, we provide a new MEG dataset of 300 simple sentences in the active and passive voice. In future, we plan to extend our analysis by: (1) source localising the neural data to gain fine-grained spatial resolution in the analysis, and (2) collecting more data to perform additional analysis pertaining to brain behaviour during question-answering.

## 7 Broader Impact

Cognitive studies for natural language understanding, endeavour to discover brain mechanisms for language understanding. The non-invasive nature of MEG/fMRI/PET/EEG studies makes a lot of data available for the same. In the past, such an experiment required careful choice of stimuli to discover the impact of a condition of interest on the brain. In contrast, Machine learning enables naturalistic experiments by discovering discriminative information about a condition of interest. Deep Learning has recently been applied to this kind of study by relating brain and language understanding machines. The core principle of this match is to be able to detect information flow in the brain using better and more compact representations of language using deep learning as compared to the alternative one-hot vectors or discreet labels. In this paper, we propose a methodological approach to detect representations of words from the past in the current context composition in the brain. Enabling a better understanding of language understanding in the brain which should hopefully inform better design of AI models for language understanding tasks. However, the findings of this study are limited to the native English language speakers. The small sample size of both the stimulus and the subjects also leads to bias in the findings of such studies.

Societally, understanding brain representations is considered key to discovering true intelligence and to improve machine learning methods to reach the accuracy levels of a human. However, we also have to be careful to avoid any biases in the development of such algorithms. The data and algorithms should be general and include information about minority population etc. In the context of our study, this would mean studying the language comprehension for a variety of different languages to discover the true language comprehension.

As we deepen our understanding of language processing in the brain, this is likely to inspire more sophisticated Natural Language Understanding (NLU) machines. While such advanced NLU may be used for many positive uses such as drug discovery, information access, etc., the same technology may be used for nefarious acts such as spreading misinformation. It is therefore vital for the field to be vigilant of such misuse of technology, and develop guard against those.

### A Appendices

#### A.1 Chance Performance for Rank Accuracy

In the prediction task (stimulus features to brain activity), we score the prediction by ranking a list of gold candidate references by cosine similarity. If the correct candidate is at the top rank position, we assign the prediction a 100 % accuracy using the following formula:

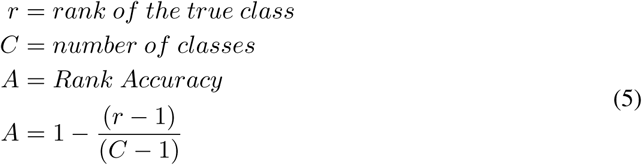

The expected value of the accuracy estimates the system performance under random prediction.

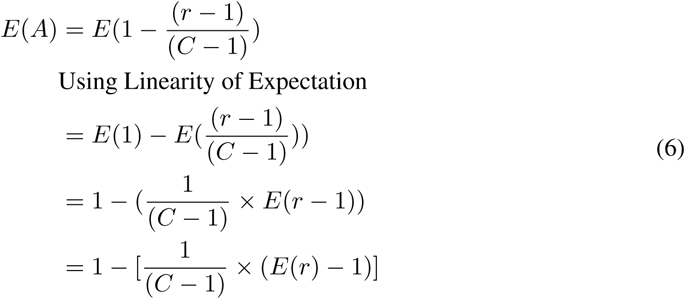

In a random prediction, the true answer is equally likely to be ranked at any of the *C* positions. Therefore, the expected value of the random rank would be 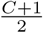.

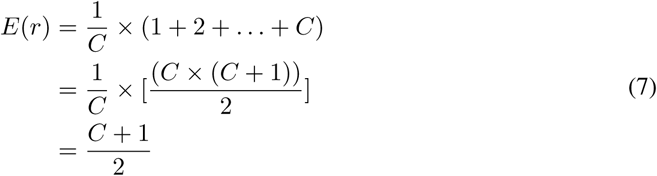

substituting Equation 7 in Equation 6

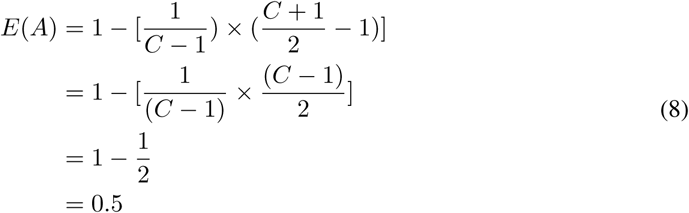

Thus proved, the chance performance of rank accuracy measure as defined in Equation 5 is 50 %.

#### A.2 SimplePassAct dataset Stimuli

The stimuli of SimplePassAct dataset is designed to be balanced for subjects and objects in the dataset. The active and passive sentences in the dataset have the following simple pattern:

*“the [noun] was [verb] by the [noun]”*

*“the [noun] [verb] the [noun]”*

Such that,

*noun ∈ [singer, baker, customer, parent, artist, author]*

*verb ∈ [encouraged, attached, answered, challenged, followed]*

We construct the sentences with all possible combination of nouns and verbs. The only condition is that a noun cannot be paired with itself in the same sentence. The total number of active and passive sentences thus formed are 300. Some example sentences are as follows:

*“the baker was answered by the parent”*

*“the parent answered the baker”*

*“the singer was answered by the parent”*

*“the parent answered the singer”*

